# Within and between sleep and cognition: associations in older community dwelling individuals

**DOI:** 10.1101/2025.10.27.684778

**Authors:** Ciro della Monica, Kiran K. G. Ravindran, Giuseppe Atzori, William Trender, Adam Hampshire, Simon S. Skene, Hana Hassanin, Victoria Revell, Derk-Jan Dijk

## Abstract

Cross sectional and interventional studies have demonstrated that sleep has a significant impact on waking brain function including alertness and cognitive performance. Few studies have assessed whether spontaneous night-to-night variation in sleep associates with variation in brain function within an individual. How this compares to the between-individual variation in sleep and cognition and their associations also remains largely unknown. These questions are of particular interest in the context of ageing because both sleep and cognitive abilities are altered in ageing. Furthermore, older people have been reported to be less sensitive to sleep loss. Here we investigated the relationship between sleep and cognition by quantifying associations between intraindividual variation in sleep and cognition as well as associations between interindividual variation in sleep and cognition in 35 cognitively intact older adults (70.8 ± 4.9 years; mean ± SD; 14 females) living in the community. Subjective and actigraphic sleep measures and daily digital assessments of cognition (nine cognitive tests; 19 variables) were obtained over a two-week period. The cognitive test battery probed a wide range of cognitive functions including reaction time, working memory, attention, and problem solving. Principal Component Analysis (PCA) identified four principal sleep components which were labelled Sleep Duration, Sleep Efficiency, Subjective Sleep Quality, and Nap-Effect. Mixed model analyses were conducted with mean and deviation-from-the-mean cognitive variables to quantify how inter- and intra-individual variation in sleep associated with inter and intra individual variation in cognition. Longer sleep duration was associated with faster reaction times in both the inter- and intra-individual analyses and with reduced errors in the inter-individual analyses. Higher sleep efficiency was associated with faster reaction times in both the intra- and inter-individual analyses. By contrast, aspects of cognition relating to learning, visual memory, verbal reasoning, and verbal fluency did not associate with sleep. The data show that in older people some aspects of waking function are sensitive to normal variation in sleep duration and efficiency which implies that interventions that target these aspects of sleep may be beneficial for waking function in ageing.

## Introduction

Sleep has a significant impact on aspects of waking function such as alertness, mood and cognition. The relationship between sleep and waking brain function is, however, complex and multifaceted and has been studied in different study designs. The methods by which sleep and cognition are assessed vary widely across these studies and may range from self-reported sleep and cognitive function to polysomnography with quantitative EEG analyses and cognitive function test batteries probing a range of cognitive domains.

Numerous studies have reported the effects of acute total sleep deprivation on waking function (e.g., (Van Dongen et al., 2003; Rusterholz et al., 2017), whereas fewer studies have used repeated partial sleep deprivation (e.g., Lo et al., 2012; Banks et al., 2024) or designs in which specific sleep stages, e.g., slow wave sleep (SWS) or rapid eye movement (REM) sleep, were manipulated (e.g., Groeger et al., 2014; Navarrete et al., 2024). These interventional studies have established the contribution of sleep to daytime alertness, sustained attention, positive affect and also provided evidence for a role of sleep in memory consolidation (Hornung et al., 2007; Koa and Lo, 2021; Groeger et al., 2022). Non-interventional studies such as cross-sectional studies have quantified associations between variation in sleep with variation in waking function across participants (e.g., della Monica et al., 2018). As an alternative to these interventional and cross-sectional designs, within-participant study designs, in which sleep and cognition are assessed daily for days or weeks, can be employed to assess the impact of night-to-night variation in sleep on day-to-day waking function within individuals.

Interventional, cross-sectional, longitudinal, and short-term within-participant designs are all valuable in understanding the role of sleep in brain function across the life span or in specific age groups such as older people. However, from an every-day-of-life perspective understanding the associations between the within-individual variation in sleep and cognition may be most relevant. This is because the range of spontaneous variation in aspects of sleep is by definition within an ecologically relevant range and understanding how within-participant variations in sleep associates with waking function may identify targets for sleep interventions that can be participant specific. Within these designs, variability itself (such as variability in sleep timing) can be considered an independent variable. Several recent studies highlighted the importance of variability in sleep timing, duration and efficiency also in relation to cognition and Alzheimer’s disease pathology (Fenton et al., 2023) and odds of developing cognitive impairment (Diem et al., 2016). Finally, these designs also allow for a comparison of the strength of the interindividual associations between sleep and cognition (i.e. trait like associations) with the magnitude of the within-participant associations.

How sleep associates with cognitive function is of particular interest in ageing because both sleep and cognition show marked changes, and interindividual variation in sleep and cognition increases as we get older. Furthermore, several studies have suggested that older people may become less affected by poor sleep or sleep loss induced by total or repeated partial sleep deprivation (Adam et al., 2006; Dijk et al., 2010; Sagaspe et al., 2012; Zitting et al., 2018).

Age-related changes in the various aspects of sleep and age-related changes in the variety of cognitive domains have been well established in cross-sectional studies with age-related reductions in slow wave sleep and processing speed being prominent examples (e.g., Mander et al., 2017). Fewer cross-sectional studies have addressed how age-related changes in sleep associate with age-related changes in cognition and whether these associations persist when age itself is controlled for as a general modifying or confounding factor. In one such cross-sectional polysomnographic study, of 206 healthy adults aged 20 to 84 years, it was confirmed that SWS changes the most with ageing (della Monica et al., 2018). In this study cognition was assessed across cognitive domains such as sustained attention, response time, motor control, working memory, and executive function and largest age-related effects were observed for aspects of ‘speed’. After controlling for age and sex, it was found that between participant variation in SWS indeed contributed to individual differences in processing speed. However, variation in REM sleep and in particular number of awakenings, which both did not show marked age-related changes, explained between participant variation in accuracy and aspects of executive function (della Monica et al., 2018). Similarly, a much larger study by Djonlagic and colleagues, which applied standard polysomnographic and advanced quantitative EEG measures but was limited to older participants, demonstrated that for macro-architectural aspects of sleep, after controlling for age and sex, increased REM sleep duration and sleep efficiency were positively associated with cognition as assessed by the Digit Symbol Substitution and Trails Making B test (Djonlagic et al., 2021). A meta-analysis of how in older people polysomnographically assessed sleep relates to cognition identified REM sleep and sleep continuity as positive predictors (Qin et al., 2023) whereas SWS was not a significant contributor. Studies in which sleep assessment was based on actigraphy have also highlighted the role of sleep continuity (e.g. Owusu et al., 2023).

Few studies have investigated how associations between night-to-night variation in sleep associates with day-to-day variation in aspects of waking function in ageing. Master and colleagues (Master et al., 2023) demonstrated associations between night-to-night variation in sleep duration and mood in adolescents. Gamaldo and colleagues (2010) reported associations for both within- and between-participant variation in sleep duration and a range of cognitive functions in older African Americans but sleep duration was only based on self-report. In another recent study in older people with and without dementia (Balouch et al., 2022), it was found that night to night variation in sleep as assessed by actigraphy and sleep diary and represented by its principal components, i.e. duration, quality, continuity, and latency, associated with day-to-day variation in alertness, daily memory errors, serial subtraction, and behavioural problems. Night-to-night variation in sleep duration and in particular sleep continuity measures were the most powerful predictors of within-participant variation in daytime performance (Balouch et al., 2022). In that study within- and between-participant associations were not compared and the relevance of sleep variability itself for daytime function was not assessed.

In many of these studies only a very limited number of cognitive tests were used (e.g. DSST, Trail Making Test A and B). Digital platforms of cognitive testing offer a scalable solution that can be used longitudinally by individuals in their own home without the need to interact with researchers. Such approaches have been used successfully, repeatedly over months or years in a variety of clinical populations including people living with traumatic brain injury (Brooker et al., 2020; Hampshire et al., 2022; Peers et al., 2022; Lennon et al., 2023; Stewart et al., 2023). To date, digital cognitive testing has not been administered on a daily basis in conjunction with objective assessment of sleep in ageing.

In this study, we used daily, self-administered, digitally assessed measures of cognitive function at-home over a two-week period while concurrently assessing sleep by self-report and objectively by actigraphy. The test battery assessed a wide range of cognitive functions including reaction time, working memory, attention, problem solving, and verbal fluency. One aim of the study was to assess the feasibility and acceptability of the daily digital assessments of cognitive performance in conjunction with daily assessments of sleep. In addition, we aimed to investigate whether (and which) parameters of objective and subjective sleep associate with between participant variation in parameters of cognitive performance in older adults as well as whether night-to-night variation in objective and subjective sleep measured at-home predicts day-to-day variation in digitally assessed cognitive performance, i.e. (within-participant variation and association).

## Methods

### Participants & screening

The inclusion/exclusion criteria for participation in this protocol were designed to ensure a representative heterogeneous population for this age range. Following an initial telephone screening interview, participants attended an in-person screening visit to determine their eligibility which included measurement of height, weight and vital signs, self-reported medical history, symptom directed physical examination and completion of the Epworth Sleepiness Scale (ESS) (Johns, 1991), Pittsburgh Sleep Quality Index (PSQI) (Buysse et al., 1989), Activities of Daily Living Questionnaire (ADL) (Lawton and Brody, 1969), and International Consultation on Incontinence Questionnaire – Urinary Incontinence (ICIQ-UI) (Avery et al., 2004). Participants were deemed eligible for the study if they met defined inclusion/exclusion criteria including self-declared stable and controlled mental and physical health conditions, able to perform daily living activities independently, consuming < 29 units of alcohol per week, and being current non-smokers.

### Study Protocol

The study received a favourable opinion from the University of Surrey Ethics Committee (UEC 2019 065 FHMS, 02 August 2019) and was conducted in accordance with the Declaration of Helsinki and guided by the principles of Good Clinical Practice. Written informed consent was obtained from participants before any procedures were performed, and participants were compensated for their time and inconvenience.

The study protocol has been described extensively elsewhere (Ravindran et al., 2023a; Ravindran et al., 2023b; della Monica et al., 2024). Briefly, participants attended the Surrey Sleep Research Centre (SSRC) and were provided with a range of devices to use at home for 7 – 14 days for sleep, circadian and environmental monitoring. Relevant to this analysis, participants were requested to continually wear an Actiwatch Spectrum (AWS, Philips Respironics, UK) to monitor rest-activity patterns and environmental light exposure. Participants could only remove the device if it were going to get wet e.g., showering, swimming, or washing up. In addition, participants were requested to complete the paper Consensus Sleep Diary-M on a daily basis which includes questions relating to sleep timing, quality and duration, daytime naps, as well as alcohol and caffeine consumption (Carney et al., 2012). Each morning, approximately one hour after waking, participants were requested to complete their daily cognitive testing on an electronic tablet using the bespoke Cognitron testing platform (Hampshire et al., 2021). The daily battery took approximately 30 min to complete and included the following nine tests: 2D manipulations, choice reaction time, simple reaction time, digit span, learning curves, paired associates learning, self-ordered search, verbal reasoning, and word definitions which probed specific domains. Following this at-home monitoring period, participants attended the Research Centre for an overnight stay which included a 10-hour period in bed with a full polysomnography (PSG) recording (in accordance with the Academy of Sleep Medicine (AASM) guidelines) from which we assessed sleep apnoea (apnoea-hypopnea index, AHI).

### Data Analysis

#### Sleep Diary

The diary (Carney et al., 2012) was completed on paper and the recorded information was converted into electronic format and reviewed for completeness and integrity. Nonsensical data was eliminated from further analysis. Sleep estimates were taken directly from the answers provided by participants (e.g., time taken to fall asleep, number of awakenings) or were derived from the answers (e.g., sleep efficiency = (total sleep time – time taken to fall sleep – duration of awakenings)/attempted sleep period).

#### Actiwatch

Activity data were collected at one-minute epochs and analysed using the proprietary algorithm in the Actiware 6.0.7 software. Each epoch was assigned as either sleep or wake using the medium sensitivity threshold (40 activity counts) and sleep summary estimates were derived. In accordance with AASM guidelines, the analysis interval for the major nocturnal sleep episode for each night was set using the clock times for Attempting Sleep and Final Awakening as recorded in the sleep diary (Smith et al., 2018). The sleep estimates derived from the AWS and Sleep Diary are shown in Table 1.

**Table 1.**
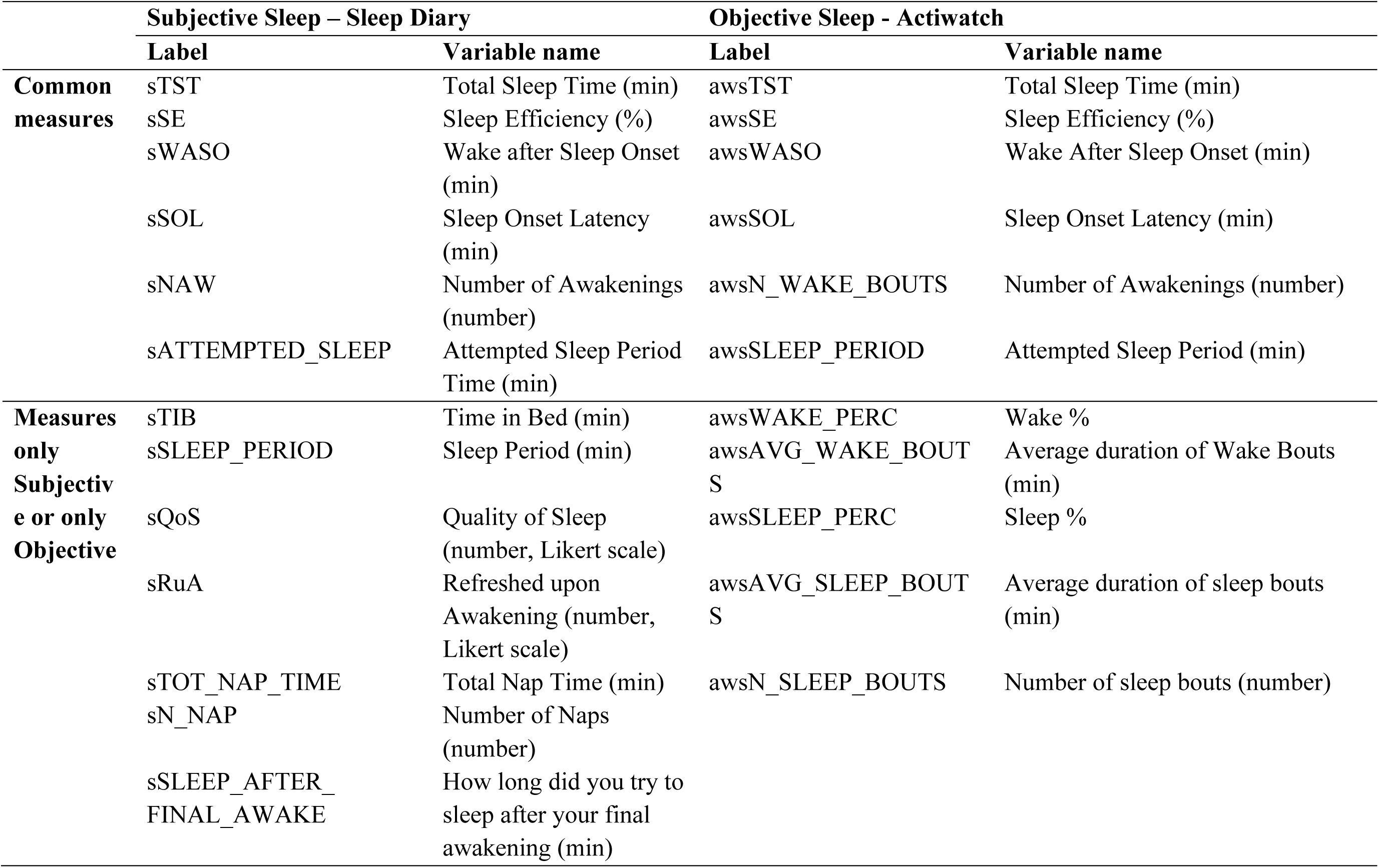
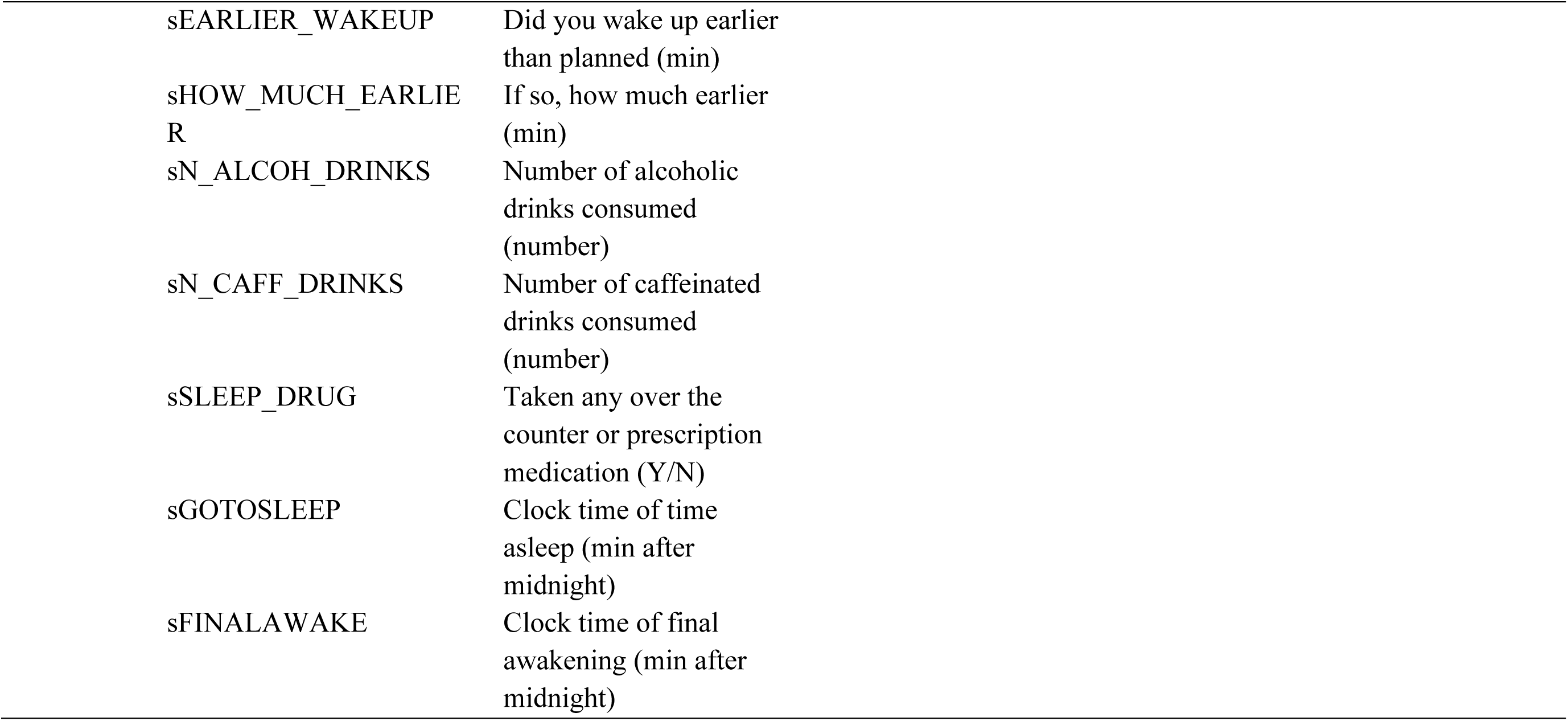
Sleep estimates derived from the Sleep Diary and Actiwatch.

#### Cognitive test battery

The data were available in .csv format for each test separately for each day and for each participant. These files were subsequently processed to extract variables relating to speed and accuracy for each test (Table 2).

**Table 2.**
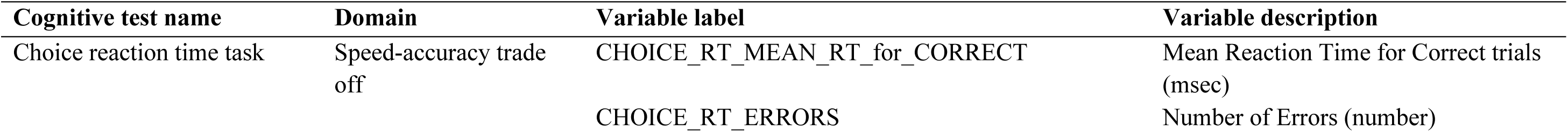
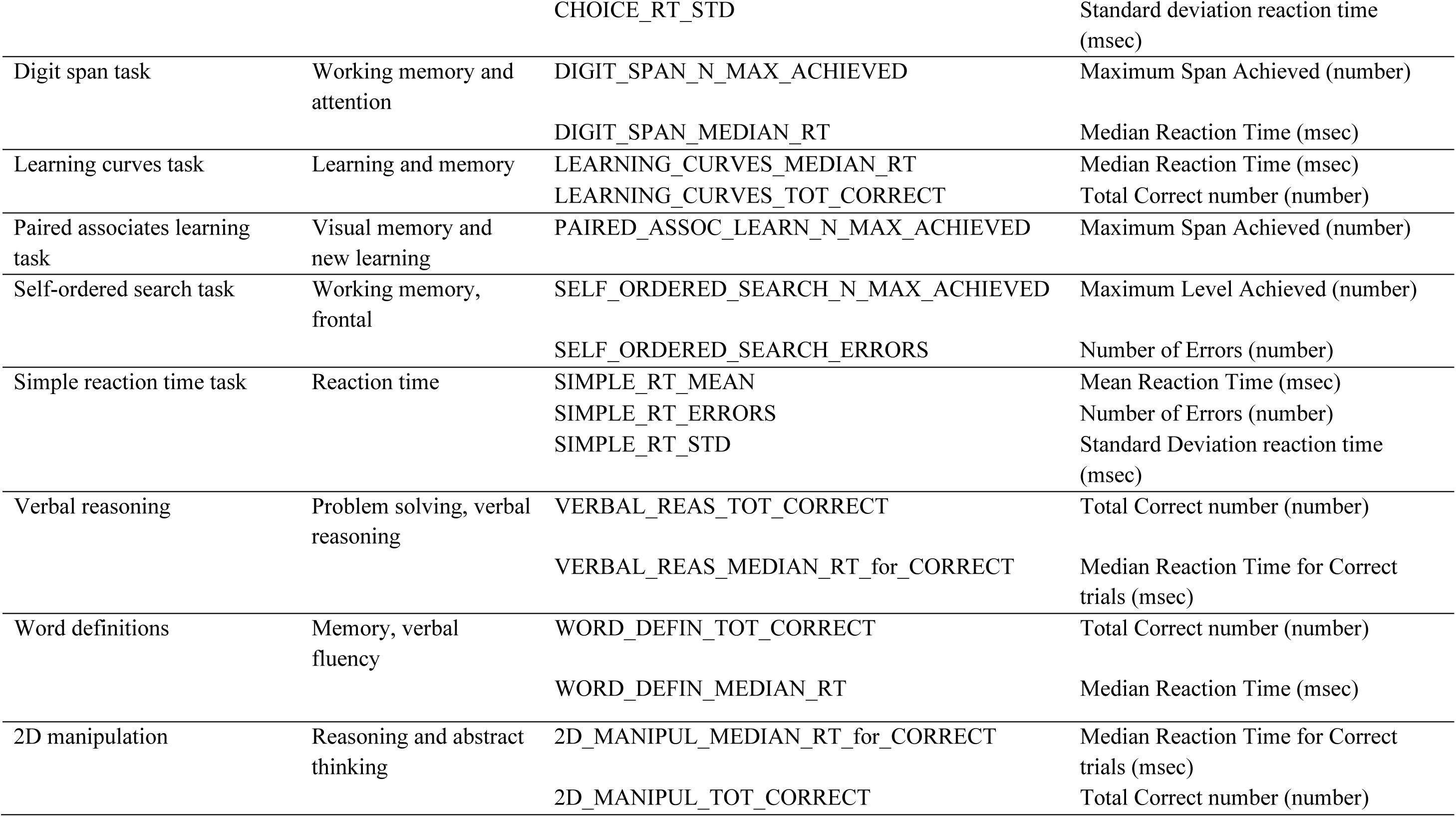
The cognitive tests used with the domain that they probe and the variables generated.

### Statistical analysis

Intraclass correlation coefficient (ICC) analysis was performed in R (v. 4.2.2, R Core Team 2022, Psych library, ICC function with two-way random effect) to determine the intra vs interindividual variation using the dependent variable (sleep or cognition) as “response” and participant ID as “subject”. An imputed dataset for subjective and objective sleep variables was created; due to the distribution of the missing data and the sample size, the approach was taken to make imputations with the mean variable value for all participants. This dataset was then subjected to Principal Component Analysis (PCA) using SAS (v9.4, SAS Institute Inc) Proc Factor, selecting the principal method followed by Varimax rotation.

A mixed model analysis was conducted in SAS using Proc Mixed where mixed models were fit for each cognitive variable, with the cognitive measure as the dependent variable, and sex, age, BMI, AHI, time-in study (to account for learning effects), and sleep component as linear fixed effects. This analysis used the non-imputed dataset as the SAS Proc Mixed function can handle missing data. Sleep components were separated into between-person (i.e. the individual’s average sleep values across the study period) and within-person levels (i.e. the deviation of the sleep values from the individual’s average). Both between- and within-person levels were simultaneously included in the model (Yap et al., 2024). Individual differences in cognitive ability were modelled as a random effect to account for repeated measures on an individual.

## Results

### Study population

Thirty-five older adults (14 females) aged 65 – 83 years (70.8 ± 4.9 years; mean ± SD) participated in the study. Participants had a PSQI score of 4.1 ± 2.1, an ESS score of 3.6 ± 2.5, an ICIQ score of 1.0 ± 1.7, an ADL score of 7.9 ± 0.2, a BMI of 26.7 ± 4.7 kg/m2 and a mini mental state examination (MMSE) score of 28.7 ± 1.4 (mean ± SD).

### Data completeness

Of 399 planned days/nights of recording at home, the following data were available for analysis: 1) AWS: 378 days (95% data availability) 2) Sleep Diary: 395 days were recorded (99% data availability), 3) Cognitron data: 379 days (95% data availability).

### Longitudinal assessment of sleep and cognition at home

The raster plots in Figure 1 provide examples of rest/activity, cognitive performance, and sleep parameters over 14 days at home for two participants using the Actiwatch and Cognitron platform. In the top panel, night-to-night variation in sleep timings and durations are evident including an increased time in bed on Saturday nights. In addition, reaction times vary from day-to-day and are fastest with increased sleep duration whereas accuracy variation does not necessarily associate with sleep duration. In the bottom panel, daytime naps are observed and there is more night-to-night variation in sleep duration. Day-to-day variation in both speed and accuracy do not obviously associate with any sleep parameter.

**Figure 1.**
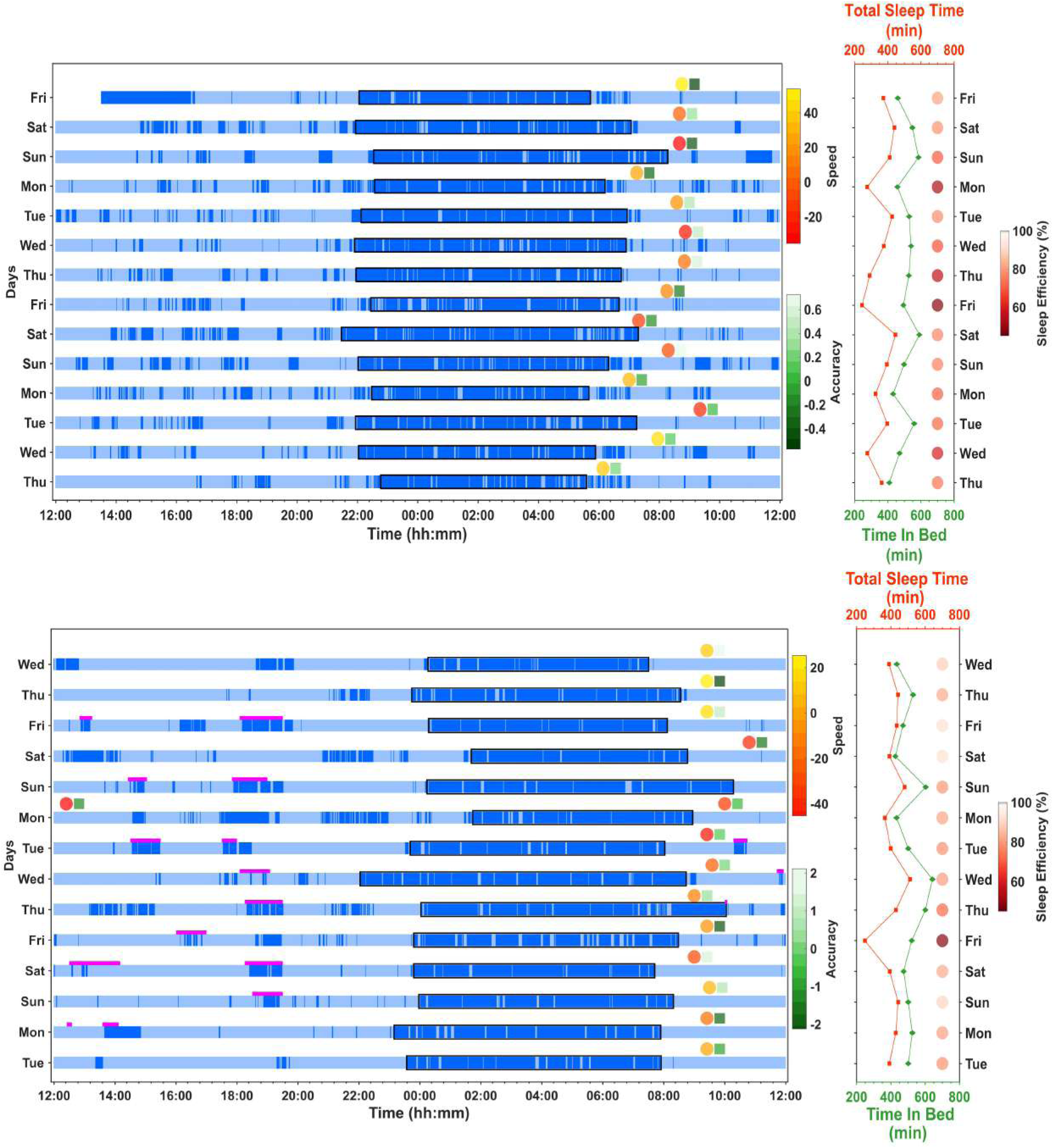
For two different participants, raster plot of rest/activity (Actiwatch) and: Top panel - Simple Reaction Time (Speed and Accuracy) and, Bottom panel - Learning Curves (Speed and Accuracy). Dark blue bars indicate periods of rest and light blue bars indicate periods of activity. The timing of the tasks is indicated by the position of the circle (speed) and square (accuracy) symbols. The spectrum indicates the accuracy or speed relative to the centred mean. For speed, positive values indicate slower reaction time and therefore, worse performance. For accuracy, in the Simple Reaction Time task positive values indicate increased errors whereas with Learning Curves positive values indicate a higher number of correct answers. Magenta bars indicate naps reported in the sleep diary (bottom panel only). The right-hand side panels indicate Time in Bed (minutes), Total Sleep Time (minutes) and Sleep Efficiency (%) for each night.

### Descriptive statistics for sleep and cognitive variables

#### Within- and between-subject variation

Examples of inter- and intra-individual variation in objective and subjective sleep measures, as well as cognitive performance measures are shown in Figure 2. The figure depicts inter and intra-individual variation and shows the variation across study days for all participants combined. Average actigraphically determined sleep efficiency varied greatly between participants, ranging from 37.32% to 95.05% but also varied considerably within participants (coefficient of variation = 19.89%, range = 57.73) (see also Supplemental Figure 1). Neither objective not subjective sleep efficiency showed a clear trend across the 14 study days. By contrast the number correct on the Verbal Reasoning task showed a clear improvement across the 14 days.

**Figure 2.**
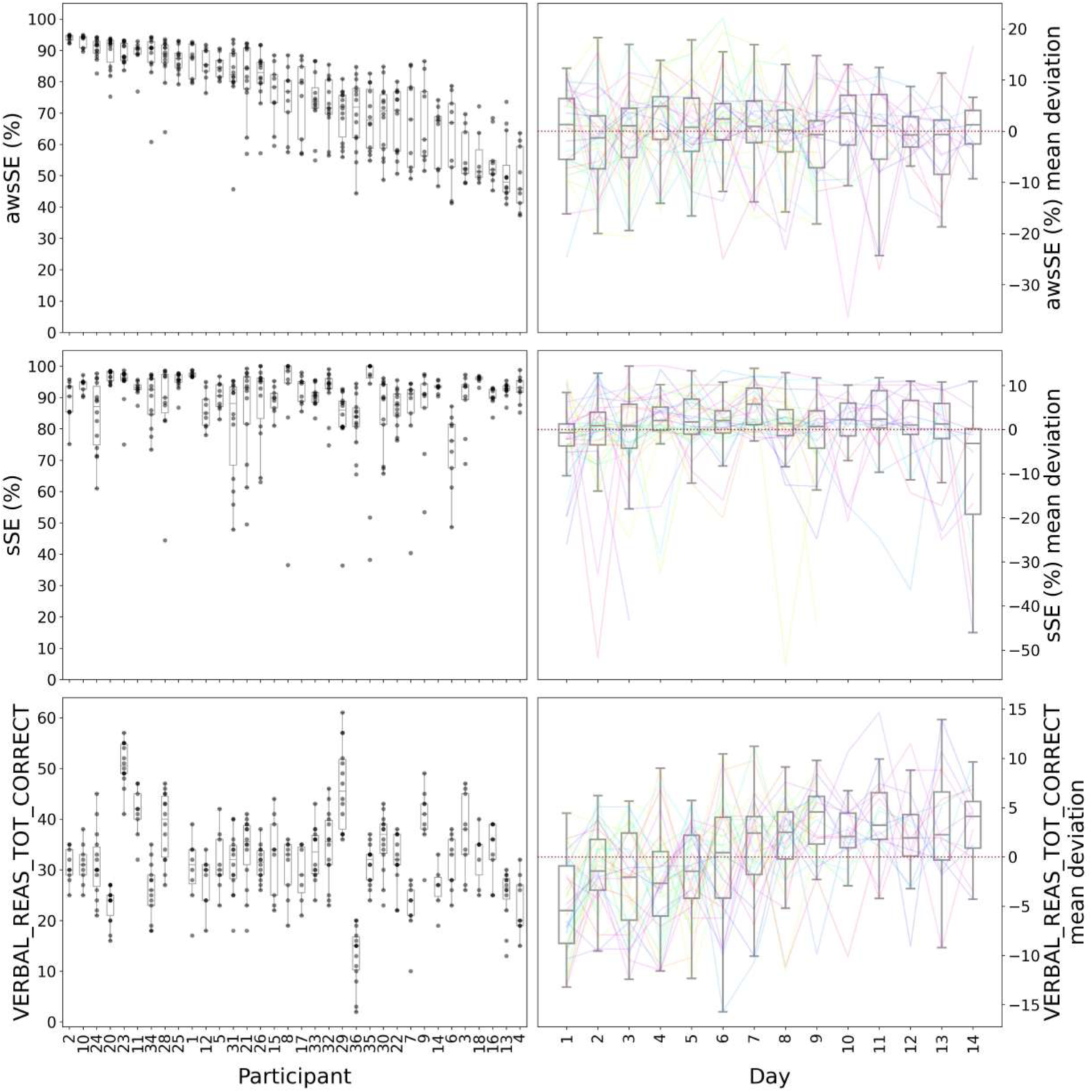
Boxplots of inter and intra-individual variation for objective sleep efficiency (awsSE), subjective sleep efficiency (sSE), and total correct answers in the verbal reasoning task (VERBAL_REAS_TOT_CORRECT) in 35 participants. In the left-hand panel, the raw data for each participant are plotted sorted by objective sleep efficiency (awsSE %). The horizontal lines represent the median, the filled circles are individual points, the boxes represent the central 50% of the data, and the vertical lines indicate the maximum/minimum values excluding outliers. In the right-hand panels the data are plotted separately for each of the 14 days. Variables were first expressed as deviation from the average in each participant and then averaged across participant for each day. Please note that performance in the verbal reasoning task improved over the course of the 14 days, whereas the sleep variables remained constant.

ICCs were computed for all of the sleep and cognitive variables (Tables 3 and 4). Values closer to one indicate that values for a participant are consistent and there is higher interindividual variation compared to intraindividual variation; values closer to zero indicate the values for a participant are highly variable and there is a higher contribution of intra-individual variation. For the sleep variables, the ICCs ranged from 0.11 to 0.86 with the lower values relating to subjective assessments of sleep onset latency (sSOL), wake after sleep onset (sWASO), and sleep efficiency (sSE). Higher ICCs were observed for objective measures including sleep efficiency (awsSE), number of awakenings (awsN_WAKE_BOUTS), and WASO (awsWASO). For the cognitive variables, the ICC ranged from 0.19 to 0.92 with the highest values (> 0.75) relating to reaction time variables (e.g., CHOICE_RT_MEAN_RT_for_CORRECT, SIMPLE_RT_MEAN, LEARNING_CURVES_MEDIAN_RT) and the lowest predominantly to accuracy (e.g. LEARNING_CURVES_TOT_CORRECT, SELF_ORDERED_SEARCH_ERRORS, PAIRED_ASSOC_LEARN_N_MAX_ACHIEVED).

**Table 3.**
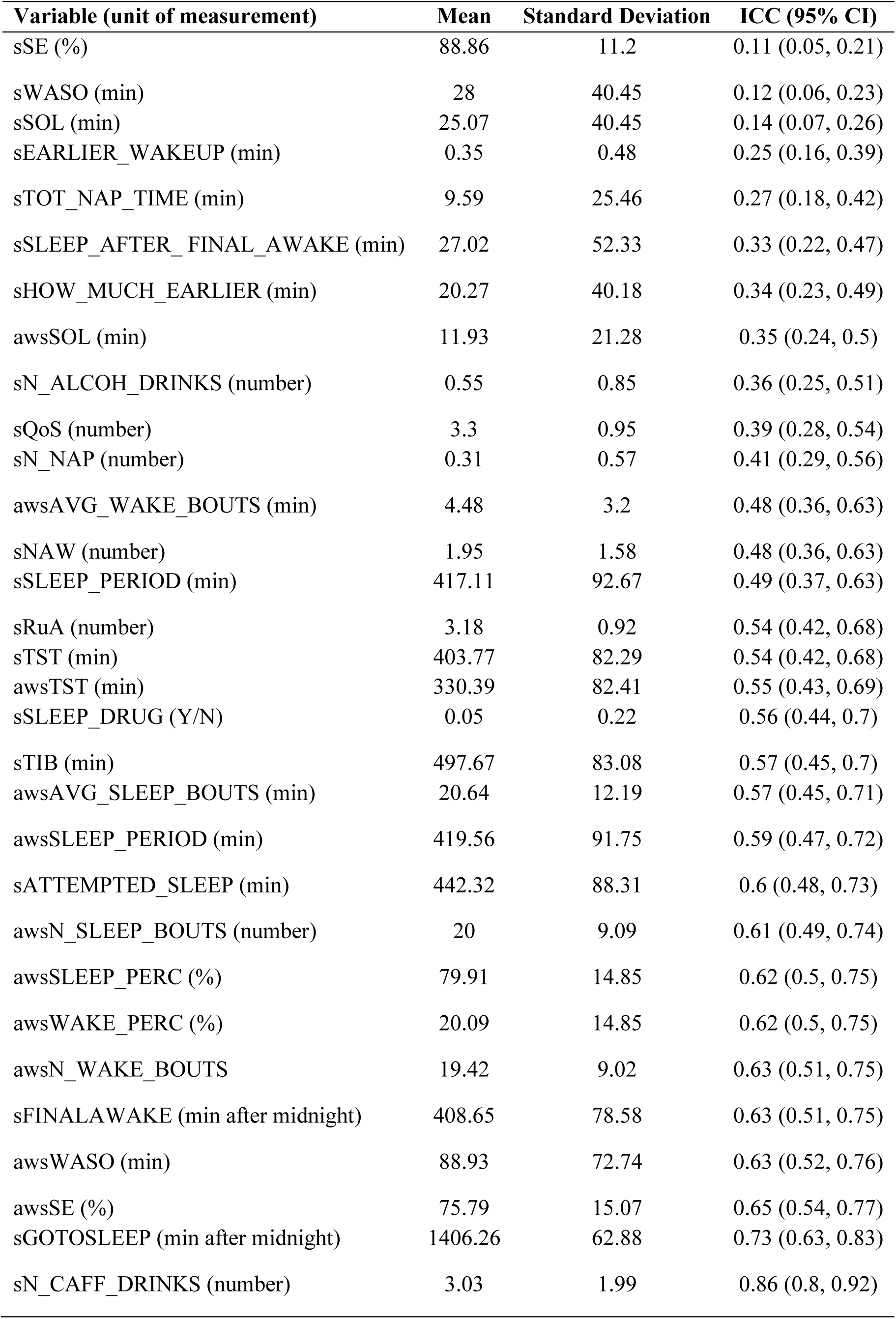
Intra-class correlations for objective and subjective sleep variables.

**Table 4.**
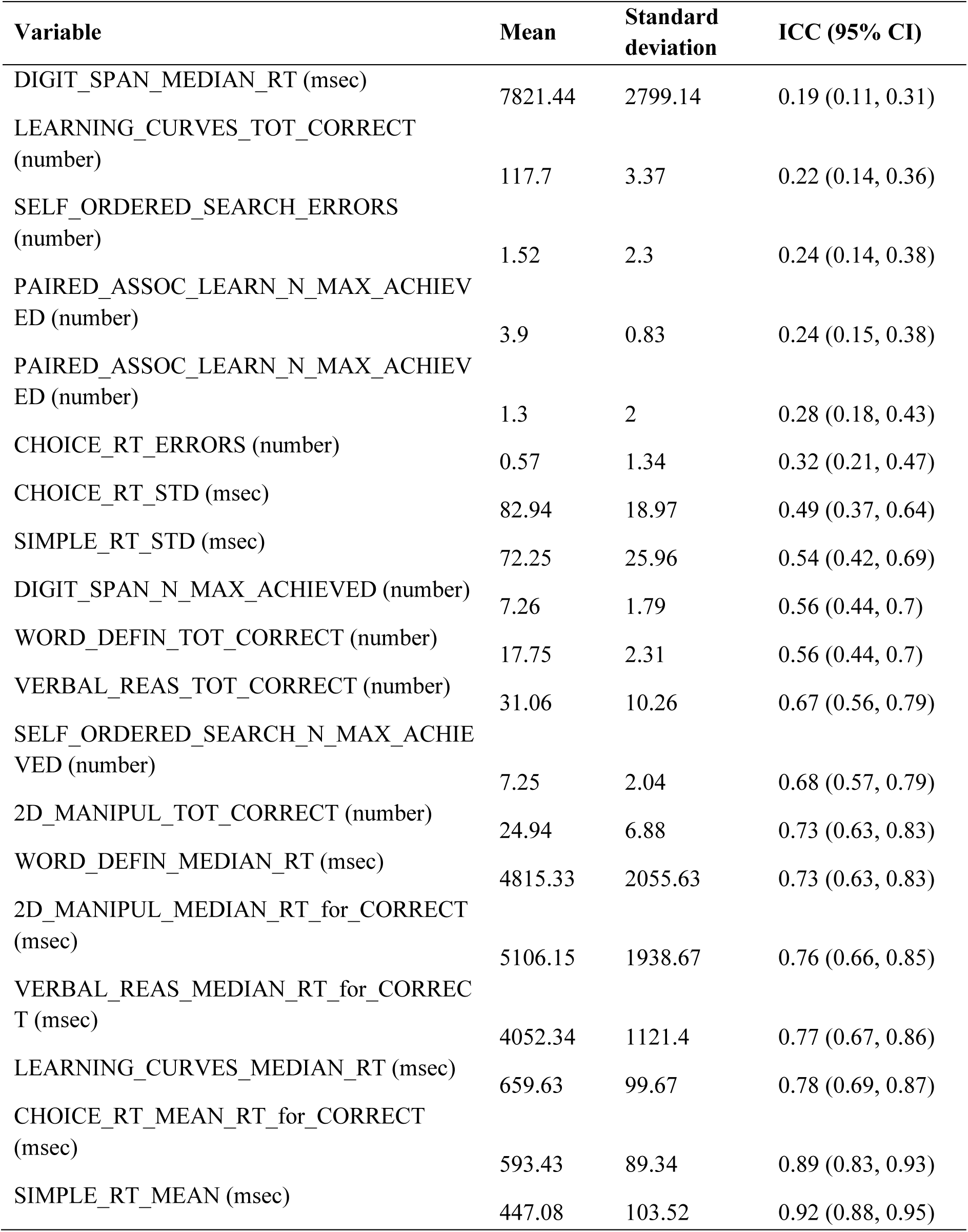
Intra-class correlations for cognitive variables.

#### Principal Component Analysis

Initial analyses of the objective and subjective sleep measures revealed that many of them were correlated and therefore, we implemented a data reduction approach, as in Balouch et al. (2022). Principal component analysis was performed including 28 of the 29 objective (actigraphy) and subjective sleep diary variables. The first four components were selected using the parallel analysis method, accounting for 54% of the total variance in the sleep variables and were interpreted as and labelled: 1) Sleep duration, 2) Sleep efficiency, 3) Subjective sleep quality, 4) Nap effect (Figure 3). The contribution of the variables ordered by sleep component is shown in Supplemental Figure 2.

**Figure 3.**
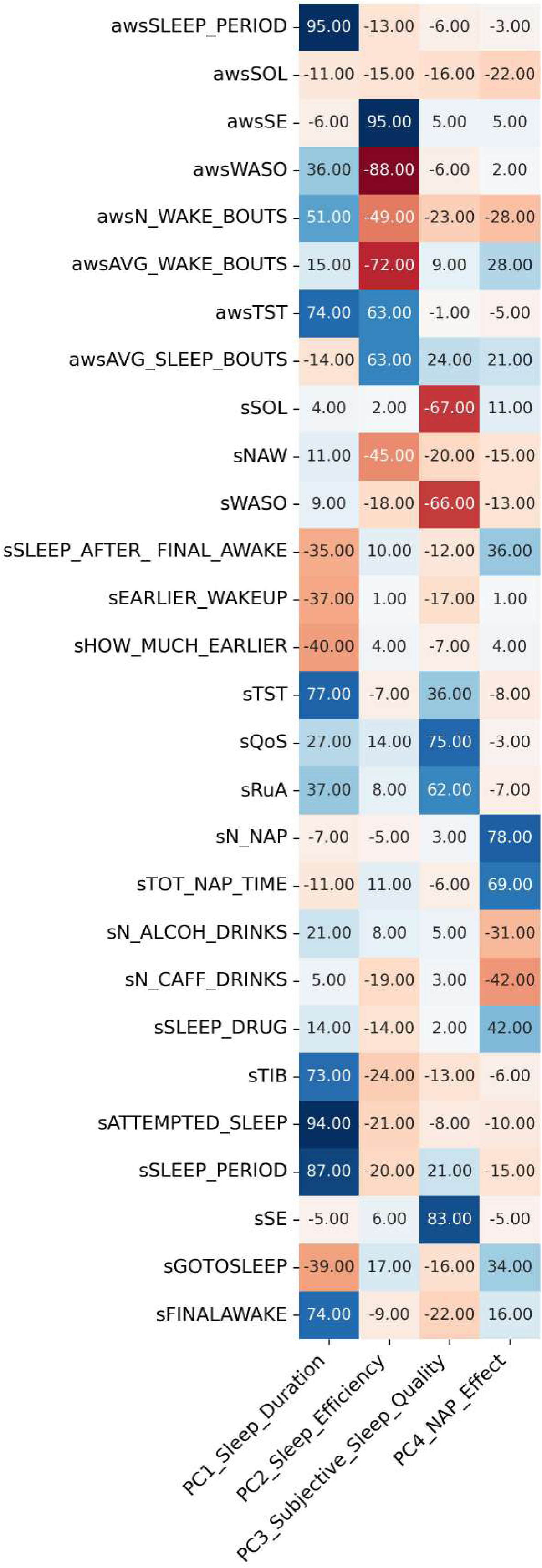
Principal component analysis of the sleep measures. Principal components analysis of the sleep measures identified four components. The strength of the contribution of the subjective and objective sleep measures to each component is indicated by colour. Blue: positive weighting. Red: negative weighting.

#### Mixed model analysis

Mixed models were conducted with each cognitive measure as the dependent variable, and age, sex, BMI, AHI, time, and sleep component as fixed effects. To assess the effects of both the intraindividual and interindividual variation of sleep on cognition, for each sleep component the individual’s average of that variable was used for interindividual effects and the deviation from the mean for each variable for intraindividual effects. There was a significant effect of Time for several variables (60 out of 76 model; each model includes a daytime cognitive variable and a principal component of sleep), which most likely reflects learning/practise effects as illustrated in Figure 2 for the total correct on the verbal reasoning task. Out of 19 cognitive variables, a significant effect of age (Sleep Component 1, 3, and 4) and AHI (Sleep Component 1, 3 and 4) was only observed for one variable (PAIRED_ASSOC_LEARN_N_MAX_ACHIEVED (the maximum score achieved on the paired associates learning task). Figure 4 provides examples of modelled associations for a female in her 70s for those associations for which the mixed model generated a significant effect of the principal component.

**Figure 4.**
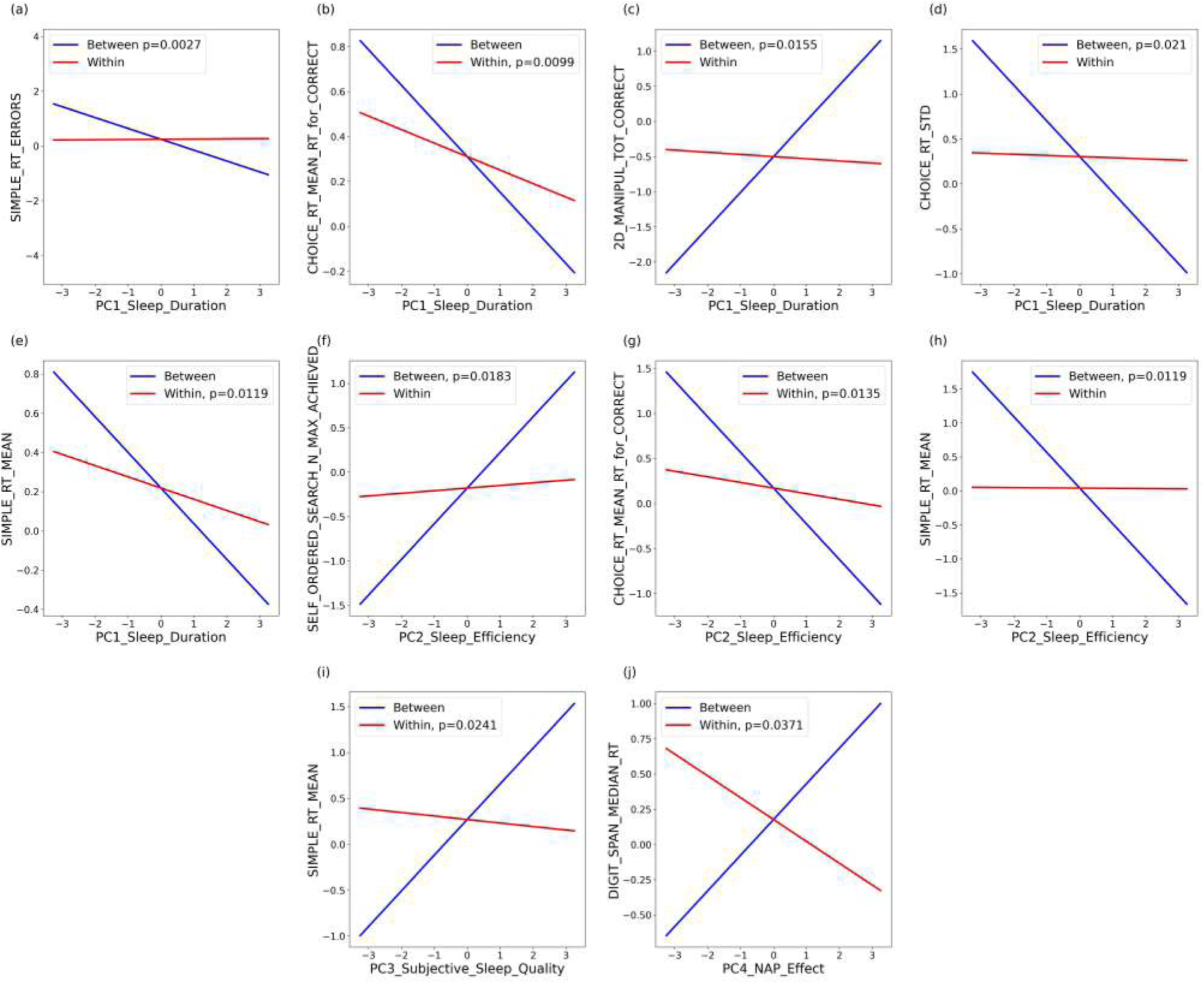
Modelled associations between daytime measures and principal components (PC) of sleep for those associations for which the mixed model generated a significant effect of the principal component. Modelled associations were created for a female in her 70s. PC1 Sleep Duration and (a) Simple Reaction Time errors; (b) Choice Reaction Time, mean for correct trials; (c) 2D Manipulation Task (total correct trials); (d) Standard deviation of Choice Reaction Time; (e) Mean of Simple Reaction Time. PC2 Sleep Efficiency and (f) Self Ordered Search Task (maximum level achieved); (g) Choice Reaction Time, mean for correct trials; (h) Mean of Simple Reaction Time. PC3 Subjective Sleep Quality and (i) Mean of Simple Reaction Time. PC4 NAP effect and (j) Median of Reaction Time for the Digit Span Task.

For the absolute (between individual) data, there was a significant effect of Sleep Duration (Sleep Component 1) on accuracy (SIMPLE_RT_ERRORS, Estimate = -0.397, F_(1,28)_ = 10.82, p = 0.0027; 2D_MANIPUL_TOT_CORRECT, Estimate = 0.5076, F_(1,28)_ = 6.65, p = 0.0155) and reaction time (CHOICE_RT_STD, Estimate = -0.3967, F_(1,28)_ = 5.98, p = 0.021). Increased sleep duration was associated with quicker response time and greater accuracy. There was a significant effect of Sleep Efficiency (Sleep Component 2) on reaction time (SIMPLE_RT_MEAN, Estimate = -0.5247, F_(1,28)_ = 7.23, p = 0.0119) and maximum achieved (SELF_ORDERED_SEARCH_N_MAX_ACHIEVED, Estimate = 0.402, F_(1,28)_ = 6.28, p = 0.0183). Greater sleep efficiency was positively associated with faster reactions and better performance.

For the deviation from the mean (within individual) data, there was a significant effect of sleep duration (Sleep Component 1) on reaction time (CHOICE_RT_MEAN_RT_for_CORRECT, Estimate = -0.0601, F_(1,340)_ = 6.73, p = 0.0099; SIMPLE_RT_MEAN, Estimate = -0.0572, F = 6.4, p = 0.0119) with longer durations associated with faster response. Reaction time was also significantly affected by Sleep Efficiency (Sleep Comp 2) (CHOICE_RT_MEAN_RT_for_CORRECT, Estimate = -0.0622, F_(1,340)_ = 6.16, p = 0.0135), Subjective Sleep Quality (Sleep Component 3) (SIMPLE_RT_MEAN, Estimate = -0.03816, F_(1,338)_ = 5.13, p = 0.0241) and Naps (Sleep Component 4) (DIGIT_SPAN_MEDIAN_RT, Estimate = -0.155, F_(1,343)_ = 4.38, p = 0.0371). Thus, greater sleep efficiency, better sleep quality, and increased naps were associated with quicker reaction times (see Supplemental Table 1).

## Discussion

Simultaneous assessment of sleep and cognition during 14 consecutive nights and days revealed that in older people both the between-participant as well as the within-participant variation in sleep associated with between- and within-participant variation in some aspects of cognition. Sleep duration and sleep efficiency were the most prominent predictors of cognition. Longer sleep duration and higher sleep efficiency were associated with shorter reaction times for both between and within associations. Variation in many other aspects of cognition were not significantly associated with sleep in either the within or between analyses of association. The relationship between sleep and cognitive function may be particularly relevant in the context of dementia to either predict its incidence, track its development, or target sleep as an intervention to slow progression. Previous studies have observed associations between cognitive decline or incident dementia with sleep duration, sleep continuity, SWS and REM sleep (Lucey et al., 2019; Sani et al., 2019; Winsky-Sommerer et al., 2019; Mander et al., 2022; Zhang et al., 2022; Li et al., 2023).

Our analyses of the intra vs intraindividual variation in sleep variables showed that subjective measures of wake in the time in bed period such as sSOL, sSE, sWASO had in general a lower ICC than objective measures sleep continuity e.g., awsSE, awsWASO. This suggests that whilst objective sleep continuity as measured by the actiwatch may be relatively stable within an individual, their recall of their sleep episode is poor and varies from night to night. Our observed ICC values for objective sleep (0.35 – 0.65) and subjective sleep (0.11 to 0.73) are comparable to those previously observed in older cohorts (e.g., Balouch et al., 2022).

Comparable analysis for the cognitive variables revealed that whilst reaction time was more consistent within an individual from day-to-day (higher ICC values) accuracy showed more daily variability (lower ICC values). PCA analysis revealed that four principal components accounted for 54% of the total variance in the sleep variables. This is lower than the comparable study by Balouch and colleagues (2022) where four components of sleep explained 75% of the variation in sleep variables. However, the identified components and their composition, i.e. the contribution of the various sleep features was very similar in the two studies.

We explored both between- and within-participant variation in sleep and cognitive performance using mixed model analysis with sleep components. Overall, when comparing participants, increased sleep duration was associated with faster reaction times and greater accuracy. Within participants, night-to-night variation in sleep duration revealed that increased duration was associated with faster reactions. Our ICC analysis revealed that accuracy showed more day-to-day variation for an individual than reaction time; the lack of significant effect of sleep components on within-participant accuracy suggests that perhaps factors other than sleep are influencing this aspect of performance.

Our observation of increased TST on performance is consistent with a study of older adults in Brazil (70.4 ± 6.8 years; mean ± SD), where verbal fluency was significantly associated with self-reported sleep duration for the past month (PSQI), with lower durations associated with poorer performance (Alves et al., 2021). However, several other studies have observed negative effects on performance of very long sleep durations. In a study of older adults (≥ 60 years), objective longer sleep time (≥ 8 hours) assessed by actigraphy for 5 – 7 days was associated with poor performance on a working memory task (single assessment) (Okuda et al., 2021). In a cross-sectional study, the relationship between self-reported sleep duration (over the past month) and cognitive function was explored in independent older adults (75.5 ± 6.4 years) (Kondo et al., 2021). Those with the longest sleep duration (≥ 9 hours) had a lower MMSE score and poorer memory than those with medium sleep duration (8-9 hours) (Kondo et al., 2021). A study in community-dwelling older women (n = 782) of varying cognitive status revealed that those with the longest TST (actigraphic) had the lowest MMSE scores and greatest odds of impaired performance particularly in verbal fluency. Higher WASO was also associated with poor recall performance and lower MMSE (Spira et al., 2017). In 1496 US older adults (≥ 60 years) from a nationally representative sample, long self-reported sleep duration was associated with poorer verbal memory, semantic fluency, working memory and processing speed as well as greater subjective cognitive problems (Low et al., 2019). Finally, a study of 2893 older Korean adults, demonstrated that in cognitively intact participants, long sleep latency and long sleep duration (PSQI) were associated with an increased risk of cognitive decline whereas a late mid-sleep time appeared to be protective (Suh et al., 2018). A recent study in 1074 older adults > 65 years (Sakal et al., 2024) observed, similar to us, improved performance with higher actigraphic sleep efficiency, but they also saw worsening performance when sleep efficiency was more variable night-to-night. In 76 adults without cognitive impairment (50-83 years), increased physical activity and actigraphic sleep duration ≥ 6 hours were assessed with improved episodic memory scores on the subsequent day; sleep duration ≥ 6 hours was also associated with faster reaction times (Bloomberg et al., 2024).

Many studies have reported a U-shaped association between sleep duration and cognition, with both shorter and longer durations associated with poor performance. Long sleep duration has been shown to be a risk factor for cognitive decline (Gildner et al., 2019). In people living with Alzheimer’s Disease (PLWA), both longer and shorter sleep associated with poor performance in a task probing attention and calculation but with fewer memory errors (Balouch et al., 2022).

We also observed that reduced sleep efficiency and reduced subjective sleep quality were associated with slower reaction times and reduced accuracy. This was the case for both between and within participants. Our observations are consistent with a study in 62 healthy older adults (≥ 60 years), where increased actigraphic sleep disruption (e.g., WASO, number of awakenings) was associated with poorer episodic memory (Yeh et al., 2021). In addition, higher sleep continuity was associated with reduced daytime sleepiness and fewer memory errors the subsequent day in PLWA (Balouch et al., 2022).

A recent meta-analysis that included 72 cross-sectional and within-participant articles explored the relationship between objective sleep (measured with PSG or actigraphy) and cognition in healthy older adults (Qin et al., 2023). The authors reported that sleep duration was not significantly associated with cognition, but measures of sleep continuity were. Better performance was associated with lower WASO, higher sleep efficiency and reduced restlessness. Sub-group analysis revealed that it was tests of memory and executive function that significantly associated with sleep. The authors suggest that the lack of relationship between sleep duration and cognitive parameters may be because the studies include TST as a linear parameter rather than a quadratic or j-shaped curve with durations <6 or >7 hours associated with worsening cognition. Indeed, a meta-analysis of 18 cross-sectional and prospective studies in older adults (> 55 years) observed that both long and short sleep were associated with poor cognitive function, in particular working memory, verbal memory and executive function (Lo et al., 2016). In older adults (65-85 years) self-reported short sleep duration (≤ 6 hours) is associated with increased b-amyloid burden and impaired memory, and long sleep duration (≥ 9 hours) was associated with worse executive function (Winer et al., 2021).

There are few existing longitudinal studies that have explored the relationship between day-to-day variation in performance and night-to-night variation in sleep. One study asked older adults (n = 121, 66 – 69 years; n = 39, 84 – 90 years) to report their sleep duration each morning and completed a numerical working memory task for seven consecutive days (Lucke et al., 2022). Between-participant associations revealed that both shorter and longer sleep durations were associated with worse performance (Lucke et al., 2022). Within-participant associations revealed that overall, night-to-night variation in sleep duration was not significantly associated with performance. However, there were individual differences in this variation and appeared that average sleep duration moderated this association. Thus, for those participants that slept one hour less than the average participant, sleeping less than usual was associated with poorer performance. For those sleeping one hour more than average, daily differences in sleep duration were not associated with performance (Lucke et al., 2022).

Recently, there has been a focus on the impact of sleep regularity on health and cognitive performance (Sletten et al., 2023). For example, an analysis of 88,975 UK Biobank participants (62 ± 8 years) identified a significant non-linear relationship between a sleep regularity index (from actigraphy) and all-cause mortality hazard with mortality rates highest amongst those with irregular sleep. (Cribb et al., 2023). In a second Biobank study (n = 60,977), higher sleep regularity (actigraphy) was associated with a 20 – 40% lower risk of all-cause mortality and was a stronger predictor of mortality risk than sleep duration (Goodman et al., 2023). In the current study we observed that variable sleep duration was associated with increased reaction time, and variable subjectively reported wake time was associated with increased errors (see Supplemental Figure 3). A previous study in older adults (≥ 60 years) also saw a significant correlation between the SD of sleep timing and number of errors (Okuda et al., 2021).

## Limitations

The main limitation of this study is that, due to the field-based nature of data collection for up to 14 days, sleep was assessed using actigraphy rather than the gold-standard approach of polysomnography. However, the ability to record data in a free-living environment is a beneficial aspect. In addition, we did not apply corrections for multiple testing as this was an exploratory study; our results are in accordance with previously published literature.

## Conclusions

In conclusion, we have demonstrated the feasibility and acceptability of conducting remote, longitudinal assessments of sleep and cognition in community-dwelling older adults. We have identified that measures of both sleep duration and sleep efficiency associate with measures of performance, including speed and accuracy, and can explain both inter- and intra-individual variation. However, aspects of cognition relating to learning, visual memory, verbal reasoning, and verbal fluency did not associate with sleep. Our approach offers an opportunity to conduct long-term (days to weeks to months) assessments of sleep and daytime function in different clinical populations, including those living with dementia, to track disease progression/symptom presentation, and explore mechanistic relationships between sleep and disease. The sensitivity of some aspects of daytime function to spontaneous variation in aspects of sleep implies that targeting those sleep aspects such as sleep continuity may lead to improvement in daytime function.

## Author contributions

CdM conducted the data analysis and drafted the manuscript. DJD conceived the study, supervised the analysis, and contributed to writing of the manuscript. CdM, GA, VR, and HH contributed to the design of the study, were responsible for participant recruitment and screening, and study conduct. The devices were setup and datasets were downloaded, curated, and plotted by CdM, GA, KR, and VR. The Cognitron test battery was designed by AH and implemented for this study by WT. SS advised on the data analysis. All the authors contributed to writing the manuscript and approved the final version of the manuscript.

## Acknowledgements

We would like to thank the members of the Surrey Sleep Research Centre, UKDRI CR&T Centre, and Surrey Clinical Research Facility for their contribution to this work, in particular Daniel Barrett, Damion Lambert, Marta Messina, and Valentina Giunchiglia.

## Data availability

The data used in this study are available from the author Ciro della Monica upon reasonable request. Contact.

---

**Supplemental figure 1:** Intraindividual variability for: (A) objective sleep measures, (B) subjective sleep measures, (C) sleep principal components, and (D) cognitive performance measures. Variability is expressed as coefficient of variation (CV) computed as the average of CVs per participant. Error bars indicate standard deviations.

**Supplemental figure 2.** Principal component analysis of the sleep measures. Principal components analysis of the sleep measures identified four components. The strength of the contribution of the subjective and objective sleep measures to each component is indicated by colour. Blue: positive weighting. Red: negative weighting. The variables are ordered by the strength of their contribution (absolute value) to each component.

**Supplemental figure 3.** Heatmap of Pearson correlation between mean cognitive performance variables and standard deviation of objective and subjective measures of sleep regularity.

