## Supplementary figures and images for "Within and between sleep and cognition: associations in older community dwelling individuals"

### Image 1.TIFF

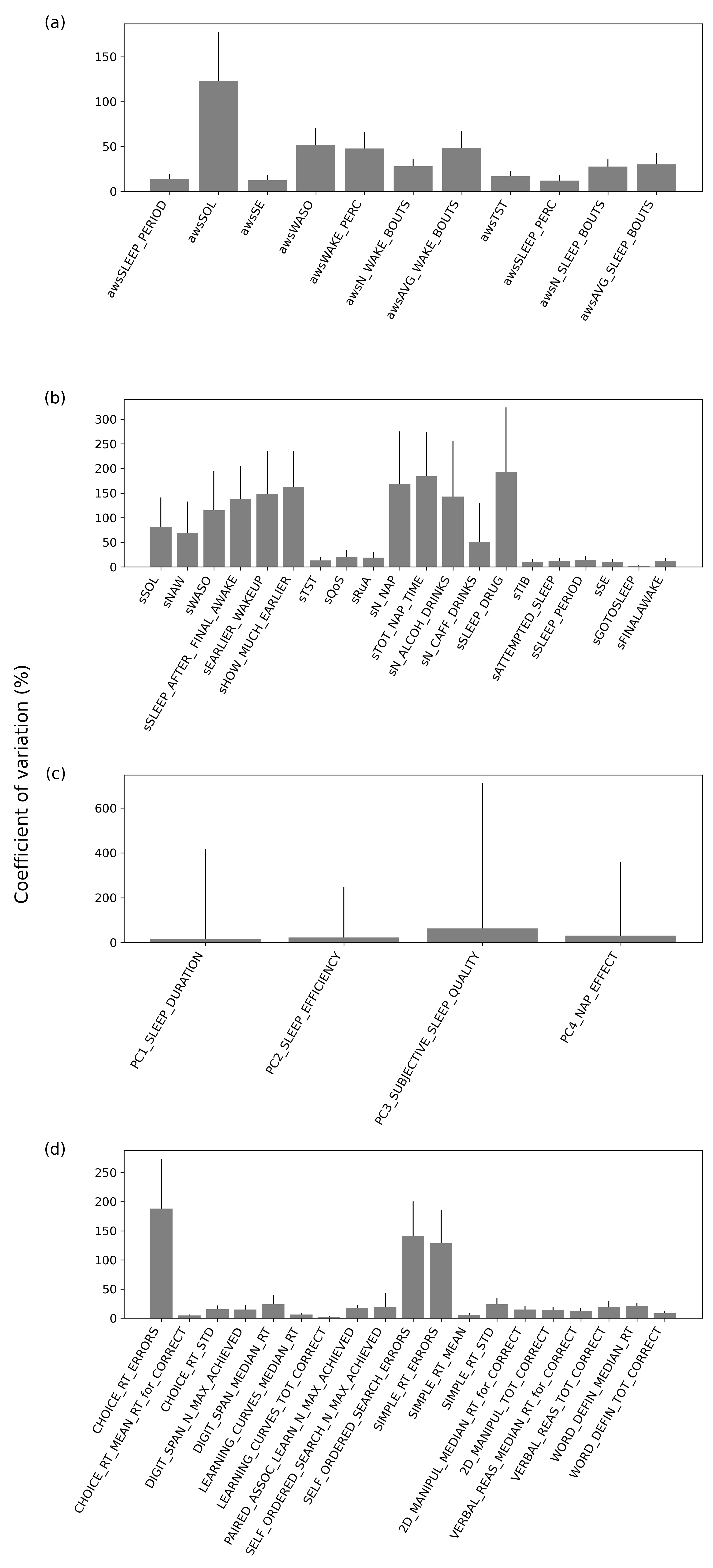

### Image 2.TIFF

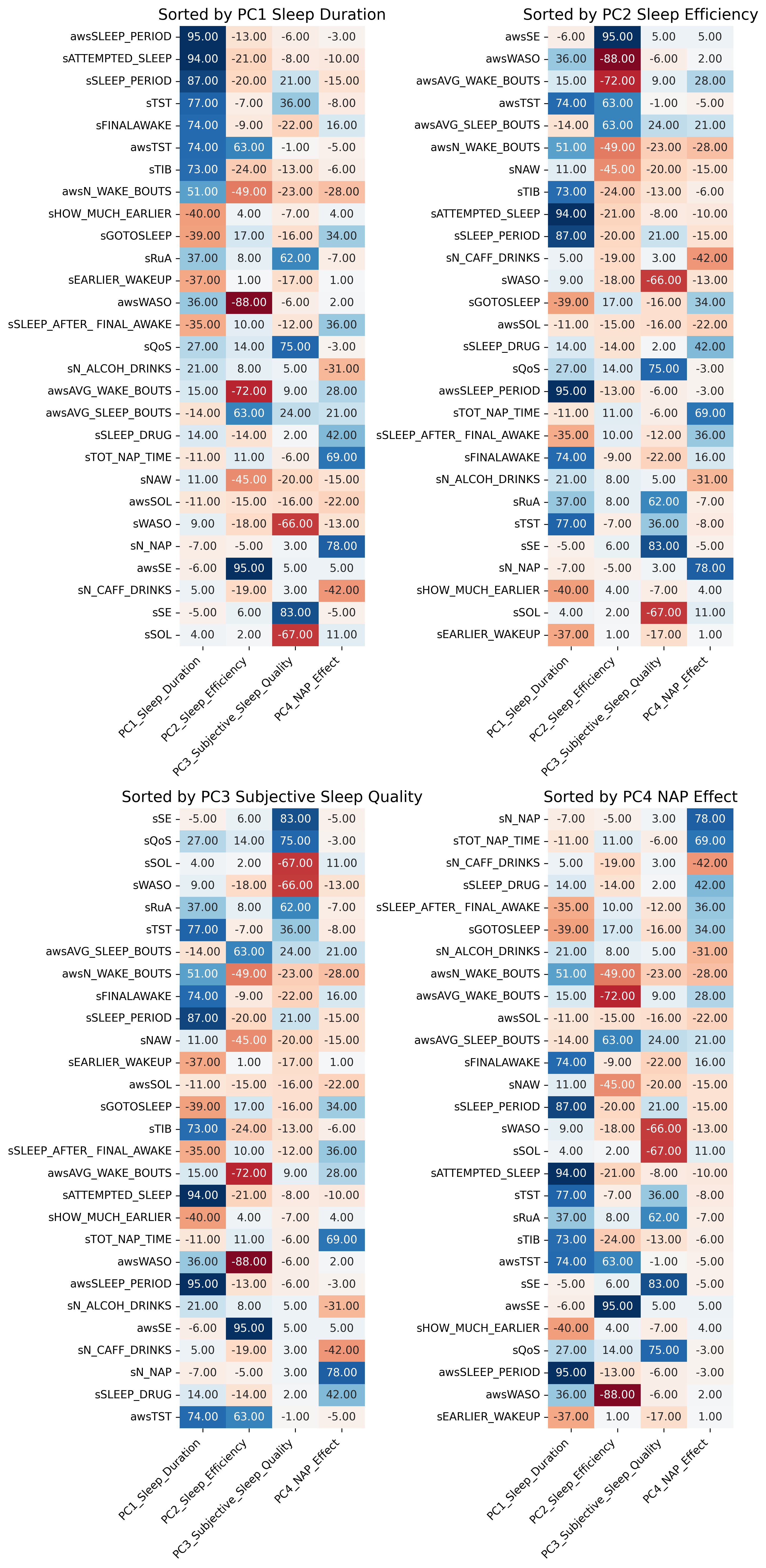

### Image 3.TIFF

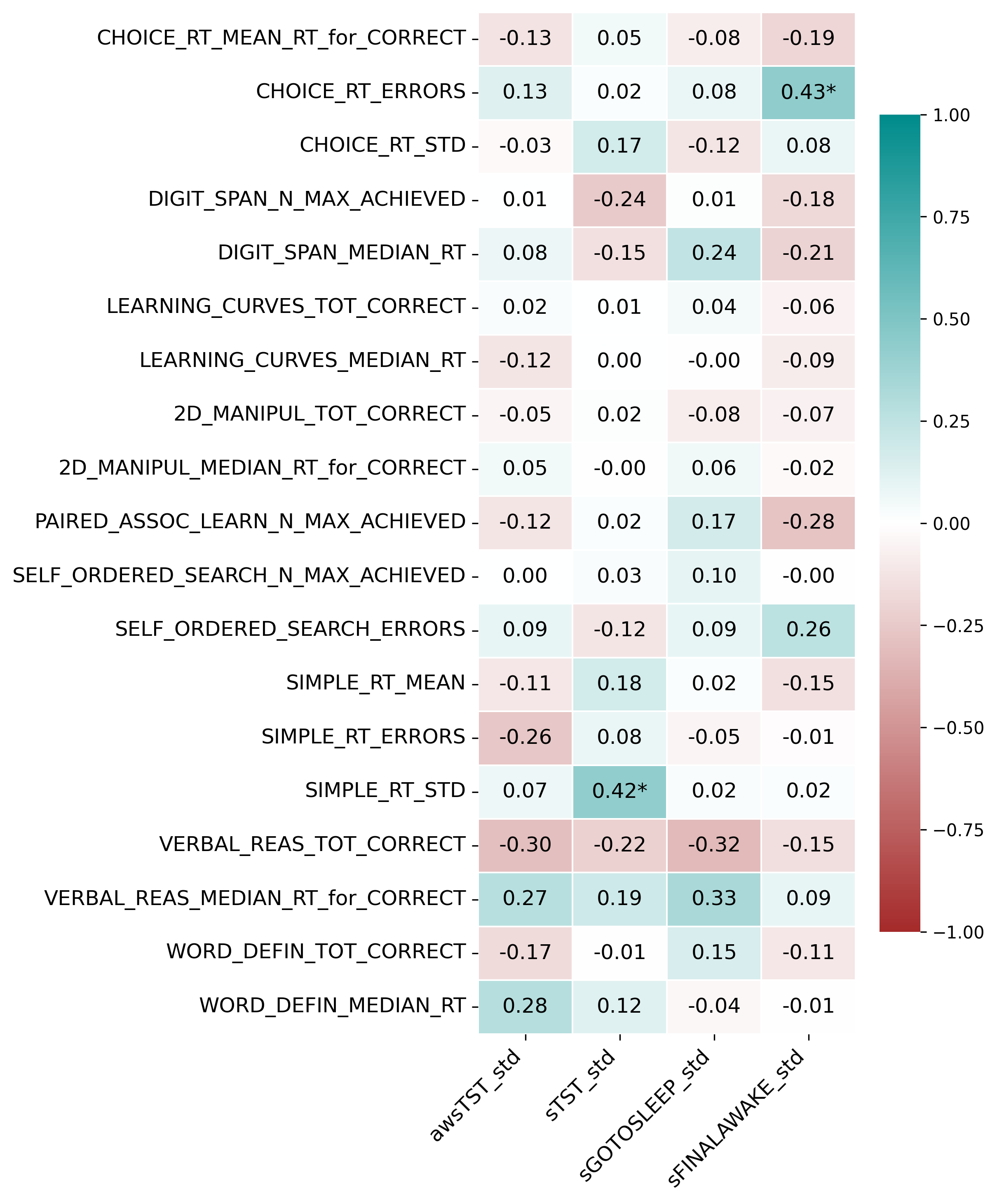
